## Supplementary figures and figure legends for "Genome architecture shapes evolutionary adaptation to DNA replication stress"

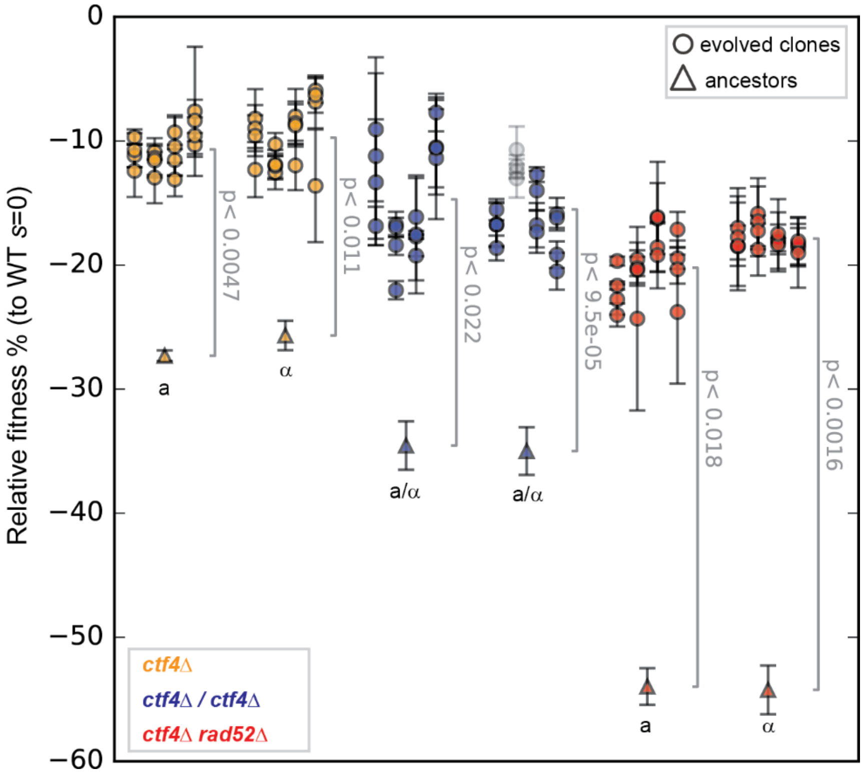


**Figure 1-S1: Fitness of the evolved clones.** Fitness of 96 clones isolated from the 24 evolved populations (4 clones for population, distributed on the same vertical line), relative to haploid or diploid WT cells ($s$ = 0). Fitness data of haploid strains (orange) is from [12]. Semi-transparent grey dots represent clones that became haploid over the course of the experiment. Error bars represent standard deviations. a and α refer to the strains' mating type (*MAT* locus). a/α indicates diploid strains. The P-values reported in figures are the result of t-tests assuming unequal variances (Welch's test). The fitness values shown here are reported in Source data 1S1.

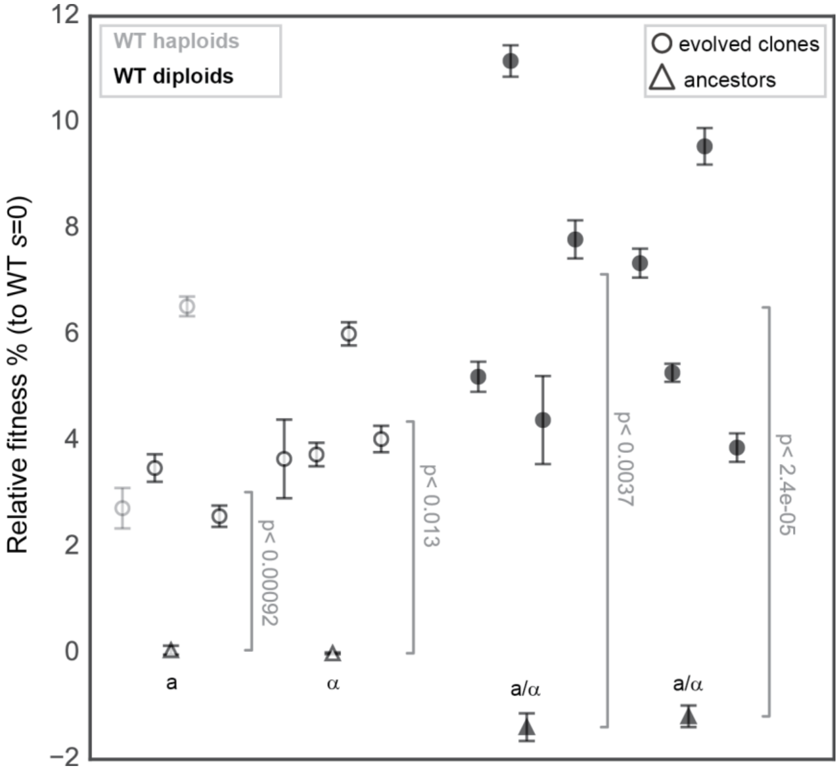


**Figure 1-S2: Fitness of the populations evolved in the absence of DNA replication stress.** Fitness of the WT haploids (white) and diploids (black) ancestors and of 16 evolved populations derived from them (four from each ancestor), relative to WT cells ($s$ = 0). Fitness data of haploid strains (white) is from [15]. Semi-transparent gray dots represent populations that changed ploidy over the course of the experiment. Error bars represent standard deviations. a and α refer to the strains' mating type (*MAT* locus). a/α indicates diploid strains. The P-values reported in figures are the result of t-tests assuming unequal variances (Welch's test). The fitness values shown here are reported in Source data 1.

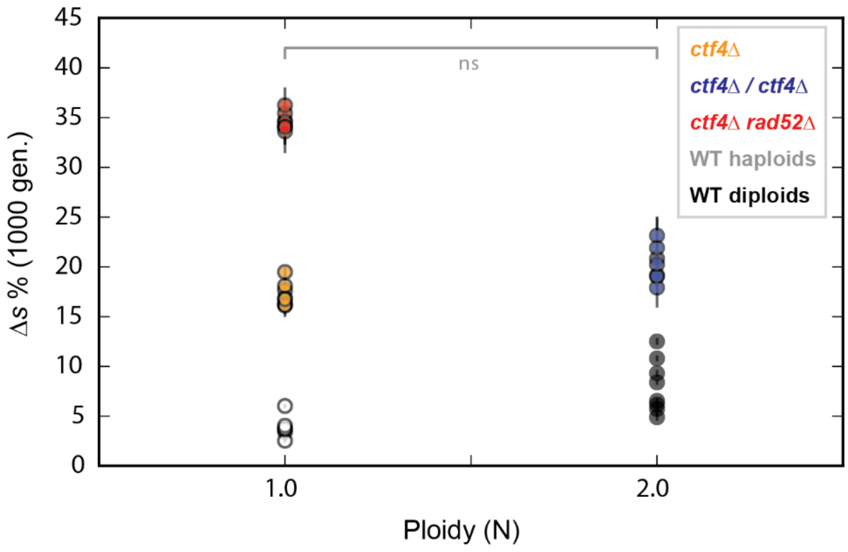


**Figure 2-S1: Fitness increase of the evolved populations relative to the ploidy of the ancestor cells.** Fitness increase over 1000 generations ($\Delta s$, measured as the difference between the populations' final fitness and the fitness of their respective ancestors that lacked Ctf4) relative to the ploidy of the ancestor cells (N). Fitness data of haploid strains (orange and white) is from [15]. Error bars represent standard deviations. The P-values reported in figures are the result of t-tests assuming unequal variances (Welch's test). The data shown here are reported or derived from Source data 1.

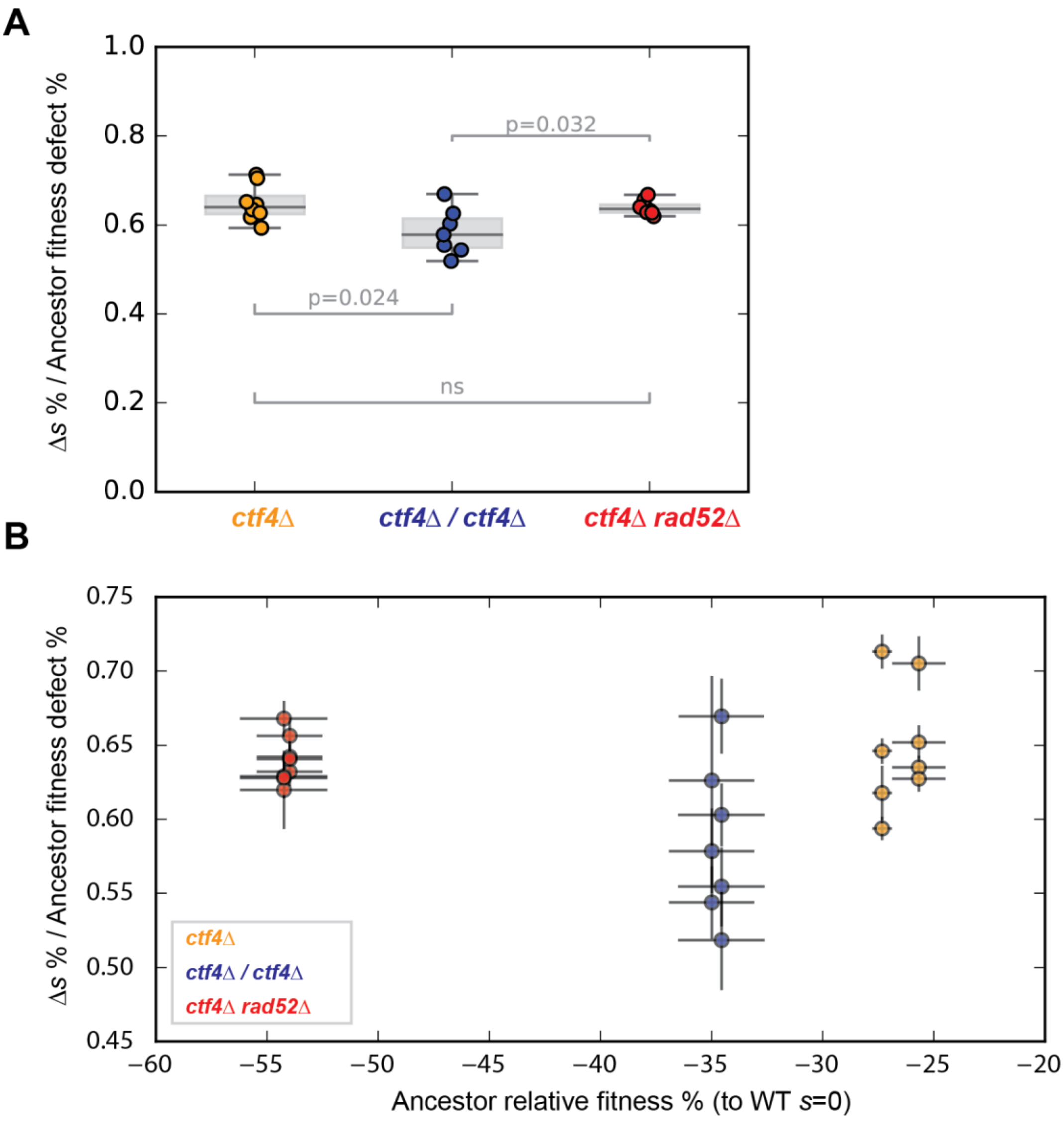


**Figure 2-S2: Fraction of the initial fitness defect recovered during the experiment. (A)** The populations' fitness increase over 1000 generations (Δ*s*) was divided by their ancestors' fitness defect (relative to strains of the same ploidy containing Ctf4) to calculate the fraction of the initial defect that was recovered during the experiment. **(B)** Fraction of the initial fitness defect recovered over the course of the experiment, relative to the ancestor's fitness. Fitness data of haploid strains (orange) is from [15]. Error bars represent standard deviations. The P-values reported in figures are the result of t-tests assuming unequal variances (Welch's test). The data shown here are reported in or derived from Source data 1.

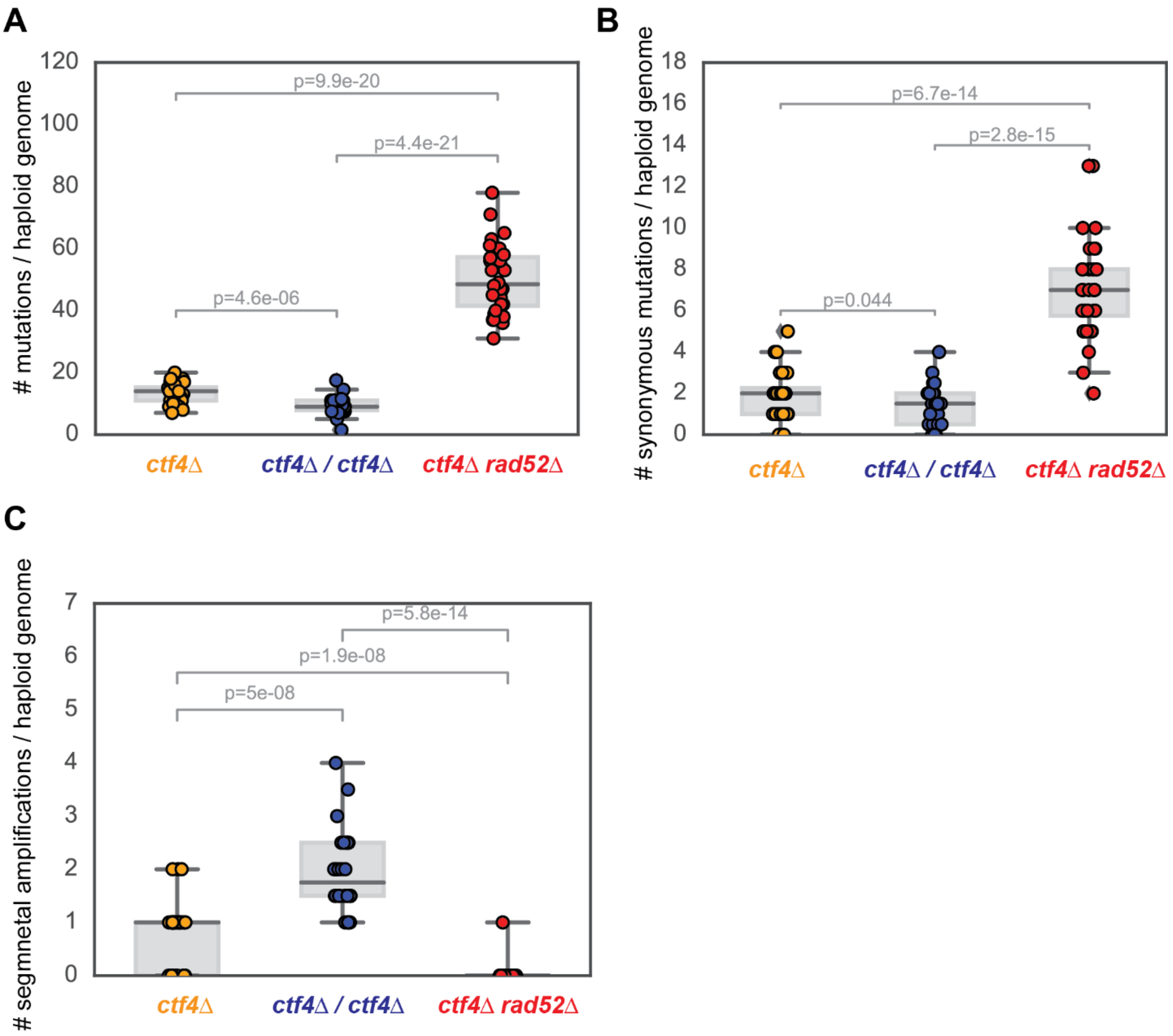


**Figure 3-S1: Number and type of mutations found in the evolved clones. (A)** Total number of mutations per haploid genome detected in the clones of each different genome architecture. **(B)** Total number of synonymous mutations per haploid genome for each different genome architecture. **(C)** Total number of segmental amplifications per haploid genome for each different genome architecture. The P-values reported in figures are the result of t-tests assuming unequal variances (Welch's test). The values shown in A and B are reported in Source data 3S1. Values shown in C are derived from Supplementary table 4.

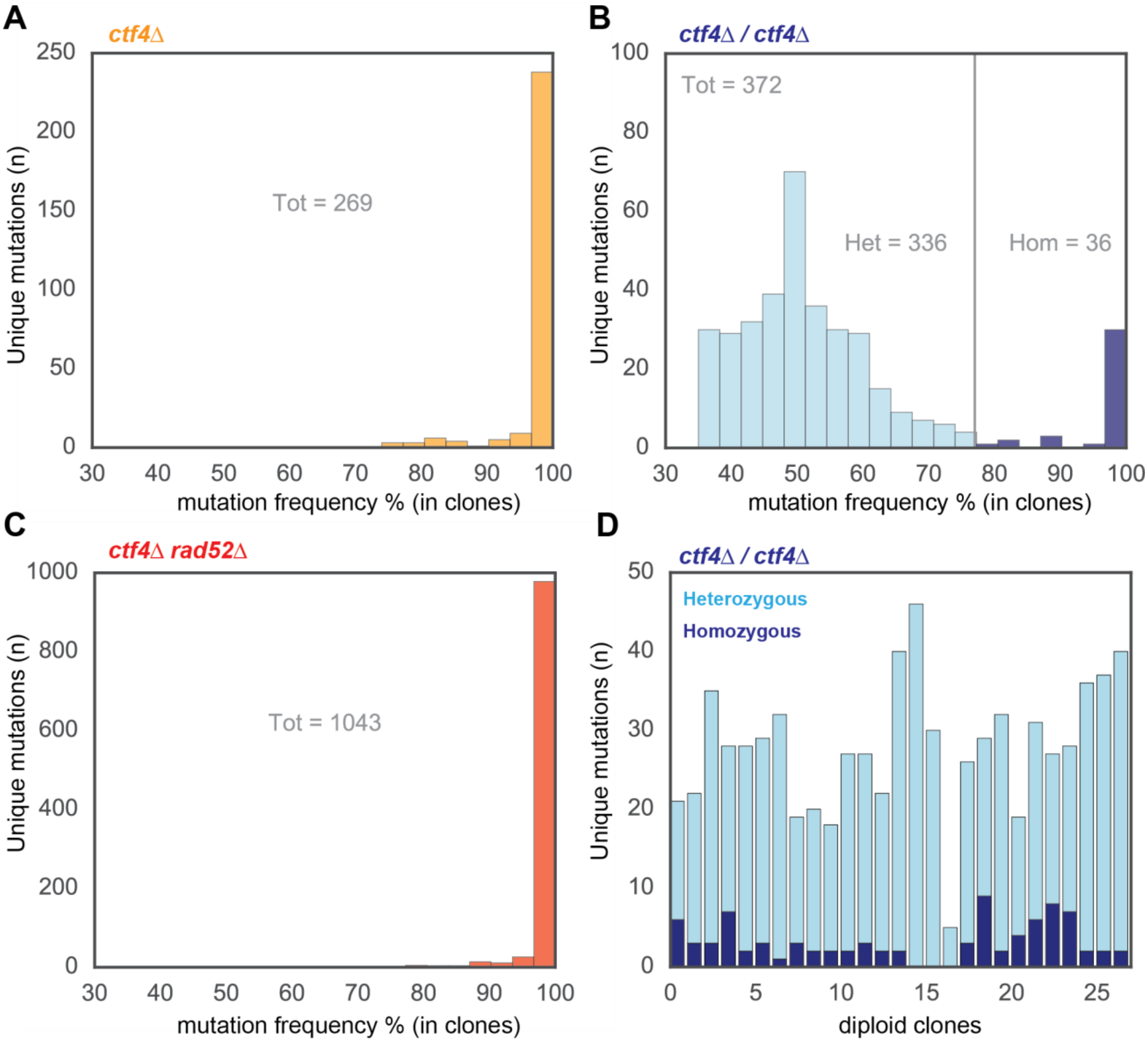


**Figure 4-S1: Frequencies of mutations detected in the evolved clones.** Frequencies of mutations in **(A)** *ctf4Δ* **(B)** *ctf4Δ/ctf4Δ* and **(C)** *ctf4Δ rad52Δ* evolved clones were obtained from the ratios of mutated and total DNA reads covering the locus. Mutations present in more than 77% of the reads were considered homozygous (Hom) in *ctf4Δ/ctf4Δ* diploids. Mutations present in less than 77% of the reads were considered heterozygous (Het). **(D)** Number of heterozygous (light blue) and homozygous mutations (dark blue) found in each individual, sequenced evolved *ctf4Δ/ctf4Δ* diploid clone. All the values shown here are derived from Supplementary table 1.

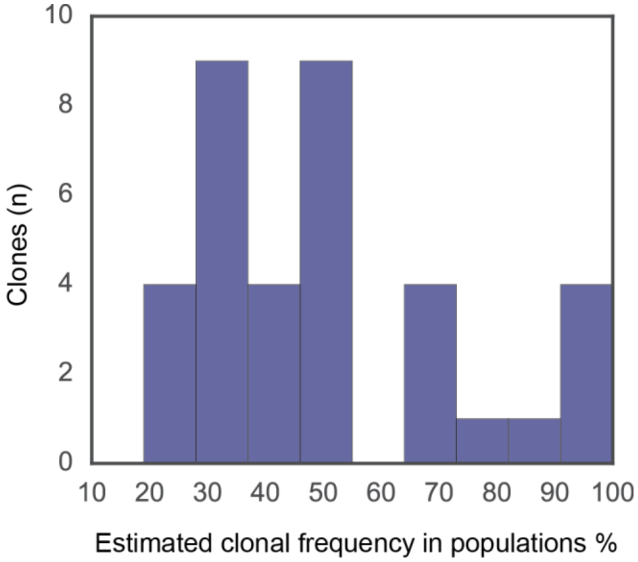


**Figure 4-S2: Estimated frequency in the evolved populations of clones carrying homozygous mutations.** Clonal frequency estimates within populations were derived from the percentage of reads from whole population sequencing that carried those mutations previously identified, in clones, as homozygous. All the values shown here are derived from Supplementary table 1.

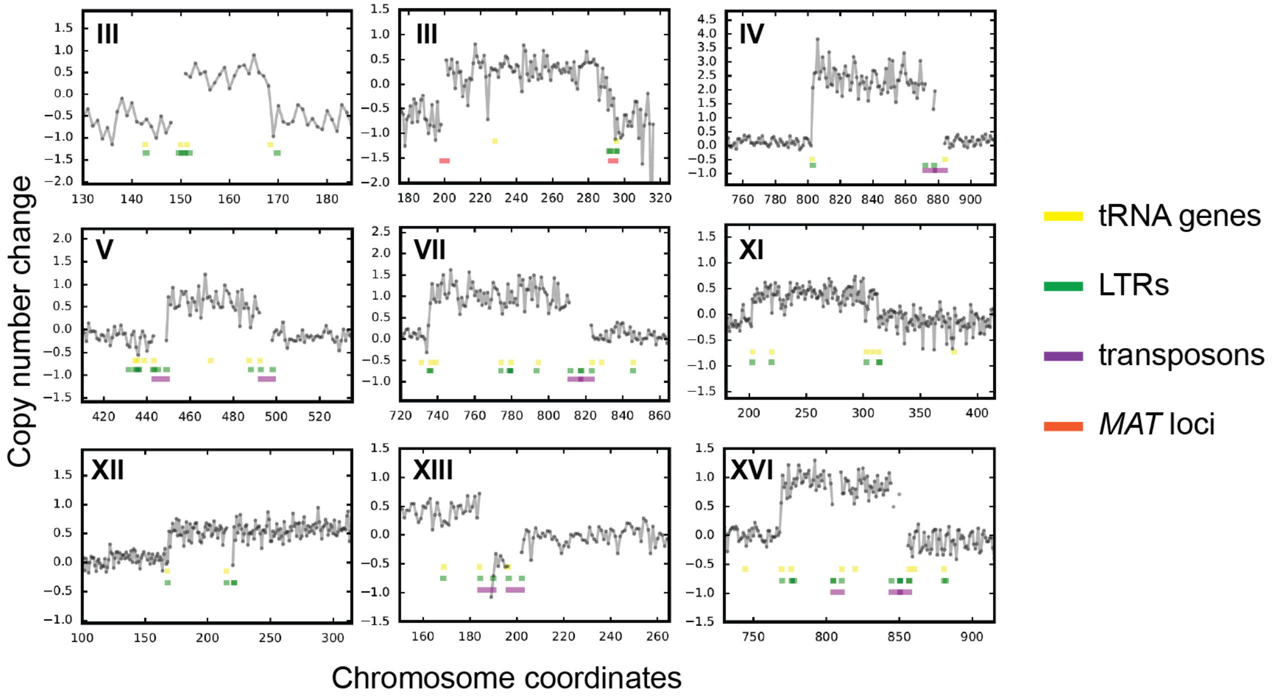


**Figure 5-S1: Segmental amplification boundaries correspond to repetitive sequences.** Magnification on the most recurrent copy number variations (CNVs) affecting different chromosomes (roman numbers). Repetitive sequences present in the surrounding chromosomal coordinates are noted in different colors: tRNA genes in yellow, Long Terminal Repeats (LTRs) in green, transposons in purple and *MAT* loci in red. Copy number change refers to the fragment's gain or loss during the evolution experiment (i.e. +1 means that one copy was gained in haploid cells, and that two copies were gained in diploid cells).

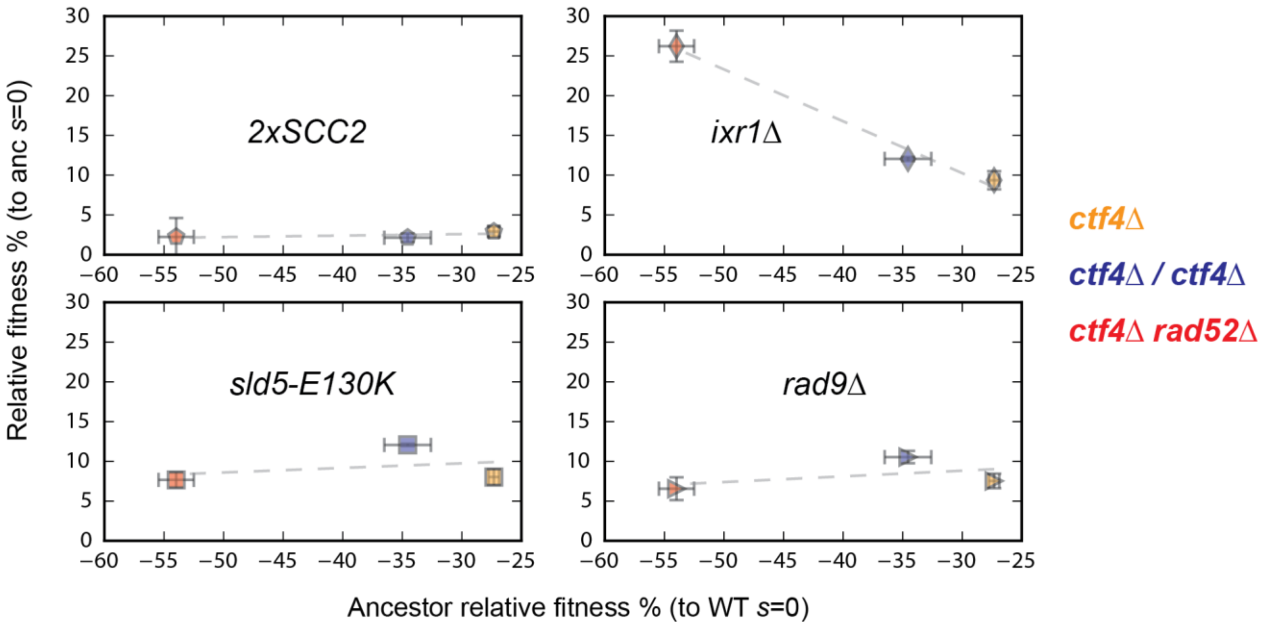


**Figure 6-S1: Limited evidence for diminishing return epistasis.** The fitness increase provided by four adaptive mutations reconstructed in different genomic architectures versus their genetic background's ancestral fitness defect. Fitness data of haploid strains (orange) is from [15]. Error bars represent standard deviations. The fitness values shown here are reported in Source data 5.
